# Deep Homology-Based Protein Contact-Map Prediction

**DOI:** 10.1101/2020.10.04.325274

**Authors:** Omer Ronen, Or Zuk

## Abstract

Prediction of Proteins’ three dimensional structure and their contact maps from their amino-acid sequences is a fundamental problem in structural computational biology. The structure and contacts shed light on protein function, enhance our basic understanding of their molecular biology and may potentially aid in drug design. In recent years we have seen significant progress in protein contact map prediction from Multiple Sequence Alignments (MSA) of the target protein and its homologous, using signals of co-evolution and applying deep learning methods.

Homology modelling is a popular and successful approach, where the structure of a protein is determined using information from known template structures of similar proteins, and has been shown to improve prediction even in cases of low sequence identity. Motivated by these observations, we developed *Periscope*, a method for homology-assisted contact map prediction using a deep convolutional network. Our method automatically integrates the co-evolutionary information from the MSA, and the physical contact information from the template structures.

We apply our method to families of CAMEO and membrane proteins, and show improved prediction accuracy compared to the MSA-only based method RaptorX. Finally, we use our method to improve the subsequent task of predicting the proteins’ three dimensional structure based on the (improved) predicted contact map, and show initial promising results in this task too - our overall accuracy is comparable to the template-based Modeller software, yet the two methods are complementary and succeed on different targets.

## 1 Introduction

Computational prediction of a protein three-dimensional structure from its sequence has seen massive progress lately due to the introduction of new deep learning models (e.g. RaptorX [17] and AlphaFold [12]). A related problem of homology modelling tackle the same prediction problem, but utilizes additional available information of known three dimensional structures of proteins that are similar in sequence to the target protein. These structures serve as templates, and can improve structure prediction beyond the performance achieved by de-novo structure prediction. Homology modelling is motivated by the observation that protein tertiary structure varies more slowly than the amino-acid sequence, hence evolutionary related proteins are likely to have similar structures. Most major recent de-novo prediction models, including the ones based on deep learning methods, also utilize evolutionary information, at the protein sequence level, including co-evolution of pairs of amino-acids [4],[12],[17]. Similarly, recent homology modelling methods use the contact maps predicted from sequence information [21] to constrain the structure prediction. However, the prediction of the contact map itself is thus far performed based on sequence only, and structural information from templates is usually used later when threading the target protein.

Here, we propose a new computational method for homology-based contact map prediction, that integrates together the sequence information from a Multiple Sequence Alignment (MSA) of a protein family, and the physical distances between amino-acid pairs for templates with known structure within this family. The integration is performed using a deep convolutional neural network. The network can accept as input alignments of different depths and different number of known template structures. Our method, called *Periscope*, utilizes both the template 3D structure information, as well as an MSA of a family of proteins, that is used to produce pairwise evolutionary couplings using methods such as CCMpred [11] and Evfold[8]. The method integrates together information from evolutionary couplings and the template structures into a deep learning architectures, and can be used when either source of information is more reliable. We evaluated the accuracy of our method in predicting a protein’s contact-map for membrane proteins and the CAMEO dataset [17]. Our method improve the accuracy of de-novo contact map prediction. Moreover, the improved contact-map can be used as constraints on energy-based methods to assist in template-based protein tertiary structure prediction, and can fold correctly proteins even when other template-based methods like Modeller [18] fail.

## 2 Methods

Consider a protein with *L* amino acids. Denote its one-hot amino-acid sequence encoding by *𝒮* ∈ ℝ^*L*×22^ (corresponding to the 20 canonical amino acids, a gap symbol and an “Xaa” symbol for an unknown amino acid), and its binary contact map by *𝒞*_*p*_ ∈ [0, 1]^*L*×*L*^, where *𝒞*_*p*_(*i, j*) denoting that the *i*-th and *j*-th residues are in contact (defined as an Euclidean distance < 8Å between the residues’ *C*_*β*_ atoms). For each protein we generate a Multiple Sequence Alignment (MSA) *ℳ* which is a family of *N* homologous proteins. For a subset *R ⊂ {*1, ..*N}* of the sequences in the family we also have known reference three dimensional structures. These structures are used to compute *r* = |*R*| ≥ 1 known distance matrices between amino-acid pairs 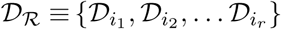, with 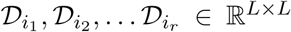 (we consider only residues aligned to the target protein, and pad with zeros distances between missing residues). The homology assisted contact-map prediction problem is defined as follows:

### Problem 1.

Given an MSA *ℳ* containing *𝒮* and a subset of known structures *𝒟*_*R*_ with *r* = |*R*| ≥ 1, predict the *contact map* of the target protein, *𝒞*. That is, find a mapping *f* with *f* (*ℳ, D*_*R*_) = *𝒞*.

We use a training set of target proteins with known contact maps to learn such a mapping 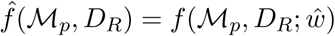, where *f* has a deep neural network architecture with parameters *w*.

### The Deep Network Architecture

We designed and implemented a neural network for predicting a protein’s contact map from sequence and structure information, shown in Figure 1(a.). The network consists of two main modules. In the first (top) HomologousNet module, we receive as input an *L* × *L* × *k* tensor representing distance matrices for template structure, and a similar tensor representing MSA-based predicted co-evolutionary matrices (we used a *L* × *L* × 2 tensor with matrices computed using CCMpred [11] and Evfold [8]). The second (bottom) module recieves as input the target and templates sequences. Each of the two modules outputs an *L* × *L* × *k* tensor, and these tensors are combined together and processed through a convolutional deep network to compute the predicted contact map.

**Figure 1:**
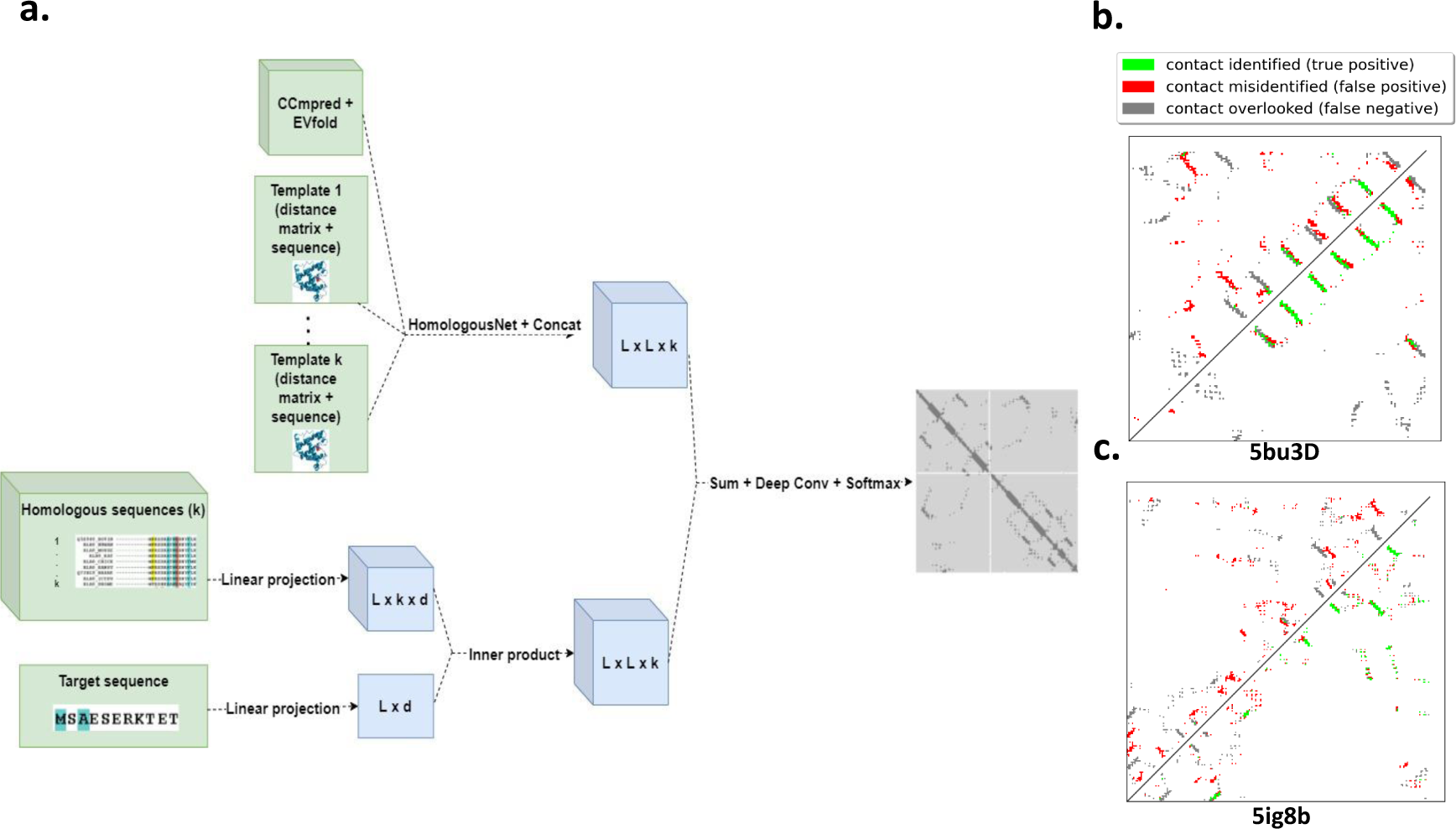
(a.) A diagram of our deep learning architecture. (b.,c.) The correct and predicted contact map for the 5bu3D (Cameo76), 5ig8B (Cameo41) proteins respectively. Top-left triangle: Modeller’s top 2*L* predictions. Bottom-right triangle: Our top 2*L* predictions. Gray squares represent missed contacts, and green squares represent identified contacts, with respect to the reference structure. Red squares represent wrong predicted contacts not appearing in the reference. Our predictions show more true positives (green) and less false positives (red), compared to Modeller.

### Details of the Training Procedure

Our test data includes the 76 hard CAMEO test proteins, 398 membrane proteins, in similar to [17], and 41 new hard CAMEO proteins. Our training set is a subset of PDB25 created in April 2020 [15]. We excluded from the training set proteins without a known structure in the alignment, long proteins (*L* > 1200), proteins with > 25% sequence identity with any test protein, and proteins with low-resolution structure (> 2.5Å), leaving us with a dataset of 9332 proteins of which 7463 (80%) were randomly selected for training and 1869 (20%) for validation. Our loss is a modified binary cross entropy between our predicted probabilities and the true zero-one contacts, averaged over all residue pairs of our training proteins. Since our classification problem is imbalanced (most amino-acid pairs don’t form a contact), we assigned a factor of 5× extra weight to positive residue pairs forming a contact. We train the model using Adam optimizer [5] for 30 epochs with learning rate *η* = 0.0001. Each training batch consisted of a single protein. To generate MSAs we ran HHblits [10] with parameters: “*-n 3 -e 1E-3 – maxfilt ∞ -neffmax 20 -nodiff -realign_max ∞* “[9] (chosen to get deep alignments). As a search library we used the uniprot20 database released on February 2016 [13]. We used SIFTS mapping [3, 14] to find solved structures among the homologous proteins (homologous that shared more than 95% sequence identity with our target were excluded). The sequences corresponding to these solved structure were re-aligned with the target structure using ClustalO [7], due to differences between the uniprot and PDB sequences. Our entire code is available at https://github.com/OmerRonen/Periscope, including a function for training our model and predicting the contact of a new example using a trained model. Additional details and documentations are available in the code repository.

## 3 Results

Figure 1(b.,c.) demonstrates our contact map predictions for two example proteins, showing that the co-evolutionary information can be used to predict contacts missed by the homology modelling method Modeller. We next evaluated systematically our method across datasets. Following [6], for a protein of length *L* we evaluate the accuracy of the top *L/k*(*k* = 10, 5, 2, 1) predicted contacts. The prediction accuracy is defined as the percentage of *native contacts* among the top *L/k predicted* contacts. We divide contacts into three groups according to the sequence distance of two residues in a contact: a contact is short-, medium- and long-range when its sequence distance falls into [6, 11], [12, 23], and ≥ 24, respectively, and report the percentage within each such group (shorter sequence distances ≤ 5 are excluded). We compare our contact accuracy to the RaptorX de-novo contact predictor [17]. We use tests set from CAMEO41, CAMEO76 and Membrane proteins (see methods). The accuracy for RaptorX is reported only in bulk, averaging over all families for each dataset. Our accuracy is evaluated on a slightly different set of proteins due to filtering (see methods). Nevertheless, the comparison shown in Table 1, is instructive - our method shows higher accuracy across most datasets and distances.

**Table 1:**
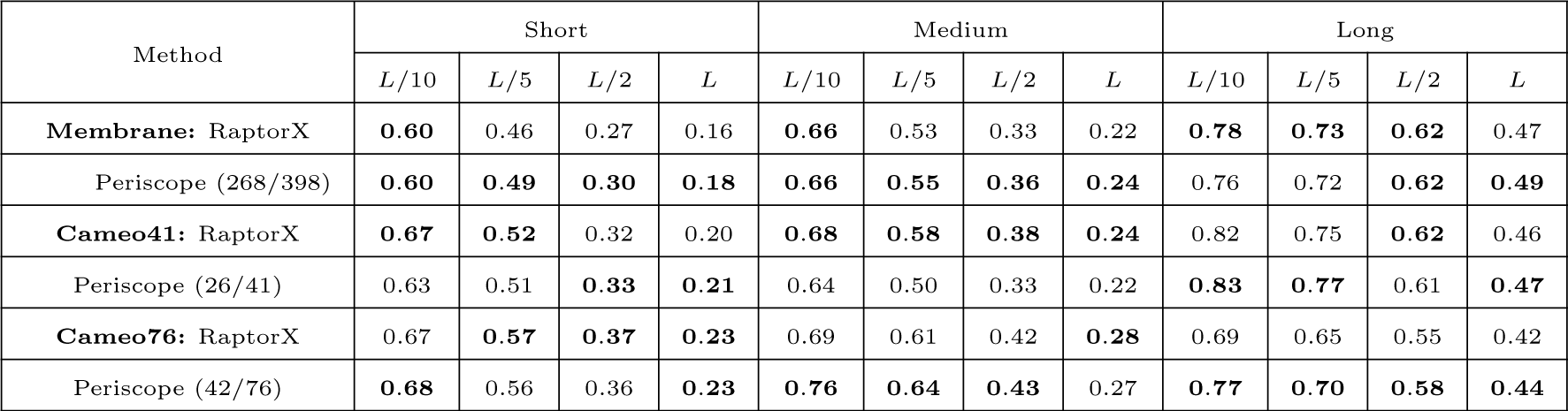
Contact prediction accuracy on membrane proteins (with 268 out of 398 proteins predicted by our method), Cameo41 proteins (26 out of 41), and Cameo76 proteins (42 out of 76).

| Method | Short |  |  |  | Medium |  |  |  | Long |  |  |  |
| --- | --- | --- | --- | --- | --- | --- | --- | --- | --- | --- | --- | --- |
| | $L/10$ | $L/5$ | $L/2$ | $L$ | $L/10$ | $L/5$ | $L/2$ | $L$ | $L/10$ | $L/5$ | $L/2$ | $L$ |
| **Membrane:** RaptorX | **0.60** | 0.46 | 0.27 | 0.16 | **0.66** | 0.53 | 0.33 | 0.22 | **0.78** | **0.73** | **0.62** | 0.47 |
| Periscope (268/398) | **0.60** | **0.49** | **0.30** | **0.18** | **0.66** | **0.55** | **0.36** | **0.24** | 0.76 | 0.72 | **0.62** | **0.49** |
| **Cameo41:** RaptorX | **0.67** | **0.52** | 0.32 | 0.20 | **0.68** | **0.58** | **0.38** | **0.24** | 0.82 | 0.75 | **0.62** | 0.46 |
| Periscope (26/41) | 0.63 | 0.51 | **0.33** | **0.21** | 0.64 | 0.50 | 0.33 | 0.22 | **0.83** | **0.77** | 0.61 | **0.47** |
| **Cameo76:** RaptorX | 0.67 | **0.57** | **0.37** | **0.23** | 0.69 | 0.61 | 0.42 | **0.28** | 0.69 | 0.65 | 0.55 | 0.42 |
| Periscope (42/76) | **0.68** | 0.56 | 0.36 | **0.23** | **0.76** | **0.64** | **0.43** | 0.27 | **0.77** | **0.70** | **0.58** | **0.44** |

### Contact-assisted protein folding

A main usage of predicted contact maps is to serve as constraints for energy-based folding algorithm in order to predict the three-dimensional structure of a protein. We used our predicted contacts, together with RaptorX-Property[16] predicted secondary structure as input to CNS-suite [2] using the CONFOLD [1] software, to predict the tertiary structure of proteins. We compared our results to Modeller, a leading template-based protein folding program using the superposition-dependent score *TMscore* [20], that measures the spatial agreement between the predicted and correct structure after alignment. As an example, the protein 5ig8B was folded with *TMScore* = 0.66 in our method, and only 0.22 in Modeller using the same templates (10 in total) (*TMScore* > 0.5 is usually considered as “correct fold” [20]). The predicted fold for the Protein 5bu3D (having 6 templates) achieved a *Tmscore* of 0.44 using our method and 0.34 using modeller. The *TMscores* for the closest template used with our targets were 0.15 for both targets. Out of the 67 proteins we predicted in the Cameo76 and Cameo41 datasets, our method achieved a *TMscore* > 0.5 in 24 proteins, while modeller achieved a *TMscore* > 0.5 in only 21 proteins. Out of our 24 good predictions, 10 were predicted poorly by modeller (*TMscore* < 0.5). This result implies that there are cases where our method can be used to fold proteins when Modeller failed, highlighting the potential for integrating templates with sequence information for prediction of contact maps and subsequent 3D folding. A more systematic comparison of the performance of these methods remains for future work.

## 4 Discussion and Future Work

In this work, we have presented, to the best of our knowledge, the first deep learning architecture for contact map prediction combining sequence information from MSAs with structural information from template homologous. While simple, our method can improve the accuracy of sequence-only contact map predictors, and may aid the correct folding of proteins when other template-based methods fail. Our framework can easily accommodate multiple improvements. For example, while we focused on predicting binary contact maps, slight modifications may enable us to predict continuous distances between amino acids [19], which can improve subsequent structure prediction. It would also be interesting to combine our method with recent threading-based methods that use the predicted contact map for homology modelling [21], or include both template and co-evolution information in complete end-to-end methods such as AlphaFold [12]. Finally, our method can be used to identify the parts of the different templates that match or disagree with the contact map of the target protein, which can be used to study the *evolution* of protein sequence and structure. With the exponential growth in the number of protein sequences, and the slower growth of experimentally verified structures, we expect homology-based contact map prediction and modelling to make ever growing impact, and aim to build upon and improve our method to handle prediction problems at a large scale.

